# Kinetics of Plasma cfDNA Predicts Clinical Response in Non-Small Cell Lung Cancer Patients

**DOI:** 10.1101/2020.08.05.238626

**Authors:** Xiaorong Zhou, Chenchen Li, Zhao Zhang, Daniel Y. Li, Jinwei Du, Ronglei Li, Effie Ho, Aiguo Zhang, Paul Okunieff, Jianwei Lu, Michael Y. Sha

## Abstract

**Background:** Tyrosine Kinases Inhibitors (TKIs), VEGF/VEGF receptor inhibitors (VEGFIs, bevacizumab) and immune checkpoint inhibitors (ICIs) have revolutionized the treatment of advanced cancer including non-small-cell lung cancer (NSCLC). Cell-free DNA (cfDNA) has been adapted as a convenient liquid biopsy that reflects characteristics of the status of both the primary and metastatic tumors, assisting in personalized medicine. This study aims to evaluate the utility of cfDNA as a prognostic biomarker and efficacy predictor of chemotherapy with or without these precision therapies in NSCLC patients.

**Methods:** Peripheral cfDNA levels were quantified in 154 patients with NSCLC before (baseline) and after (post-chemotherapy) the first target cycle of chemotherapy. These patients were divided into four subgroups receiving chemotherapy only, chemotherapy plus TKIs, chemotherapy plus VEGFIs, and chemotherapy plus immune checkpoint inhibitors (ICIs), respectively. The correlations of cfDNA with tumor burden, clinical characteristics, progression-free survival (PFS), objective response ratio (ORR), and therapy regimens were analyzed.

**Results:** Baseline cfDNA, but not post-chemotherapy, positively correlates with tumor burden. cfDNA Ratio (post-chemotherapy/baseline) well distinguished responsive individuals (CR/PR) from non-responsive patients (PD/SD). Additionally, cfDNA Ratio was found to be negatively correlated with PFS in lung adenocarcinoma (LUAD), but not in lung squamous-cell carcinoma (LUSC). LUAD patients with low cfDNA Ratio benefit significantly including prolonged PFS and improved ORR, compared with those with high cfDNA Ratio. When stratified by therapy regimen, the predictive value of cfDNA Ratio is significant in patients with chemotherapy plus VEGFIs.

**Conclusion:** The kinetics of plasma cfDNA during the chemotherapy may function as a prognostic biomarker and efficacy predictor for NSCLC patients.

## INTRODUCTION

Lung cancer is the most common cancer worldwide including both incidence (11.6% of the total cases) and mortality (18.4% of the total cancer deaths)[1]. In 2018 there was an estimated 2.1 million new cases and 1.8 million deaths, representing 1 in 5 cancer deaths[1]. The main histological categories of lung cancer are non–small cell lung cancer (NSCLC, 85% of patients) and small cell lung cancer (SCLC,15%)[2]. NSCLC consists of several subtypes, predominantly lung adenocarcinoma (LUAD, 40%), lung squamous-cell carcinoma (LUSC, 25-30%), and large-cell carcinoma (LULC, 5-10%), which are all treated in a similar manner[3]. The 3-year or 5-year overall survival (OS) of early stage (I and II) NSCLC patients undergoing resection has reached to 83% and 76% respectively[4]. Despite multiple treatment options, the 5-year OS of late stage NSCLC remains extremely low[5], with over 50% dying within one year following diagnosis[6]. Unfortunately, over one third of NSCLC cases are diagnosed at late stage (III and IV), which render them to chemotherapy, radiotherapy, targeted therapy and immunotherapy[2]. Advanced NSCLC patients are increasingly benefitting from new therapeutic advances include targeted therapies and immunotherapies[7, 8]. These precise approaches seem to produce some synergistic effects when combined with chemotherapy[9]. Thus, there is an urgent need to discover biomarkers that can assist in selecting optimal treatment, predicting response and prognostics to improve the clinical outcome of NSCLC patients.

Genotyping tumor tissue with next generation sequencing (NGS) represents an effective way to capture actionable genetic alterations as potential biomarkers in clinical oncology[10]. However, tissue biopsy may be limited due to adequacy of sample or inaccessibility for biopsy and only 25-50% of lung cancer patients have sufficient tissue for genotyping[11]. Complicating biopsy availability, the biopsy represents a single snapshot in time and is often a sample from a heterogenous location within the tumor[12], obtaining repeat specimens for genetic analysis before and after treatment is logistically difficult[13]. Therefore, liquid biopsy or blood sample becomes an alternative source and promising technology for genotyping. Increasingly, concordance has been established between liquid- and tissue-based genomic screenings[14]. Of note, some studies have suggested that liquid biopsy, specifically cell-free DNA (cfDNA), may better capture the heterogeneity of certain cancer features such as acquired resistance[15-17], and could be useful to monitor tumor burden and metastasis[18].

Emerging data have demonstrated that liquid biopsy-based biomarkers may serve as indirect indicators for NSCLC diagnosis and treatment monitoring, including circulating tumor cells (CTCs)[19, 20], circulating free tumor DNA (ctDNA)[21, 22], exosomes[23] and tumor-educated platelets (TEP)[24]. However, none of these platforms are perfect. All the above methods still have issues, making none fully satisfactory. For instance, limited CTCs detection efficiency is low, with only 32% of NSCLC patients having ≥2 CTCs using CellSearch™ (the only approved methodology by the U.S.

Food and Drug Administration)[25]. Low quantities of ctDNA in blood and sequencing artifacts may debilitate the confidence of NGS applications in detecting the actionable mutations[26]. Both CT and ctDNA are relatively time consuming and not cost-friendly for daily clinical practice. cfDNA, on the other hand, is relatively abundant and easier to quantify in circulating blood. Though the majority of cfDNA is often not of cancerous origin, preliminary studies suggest that cfDNA level and kinetics may still be used to assist in cancer diagnosis, treatment response or prognostic prediction[27-32].

However, the clinical value of cfDNA application in NSCLC has not been well-established due to inconsistent reports[33-36]. Our recent study confirmed that plasma cfDNA concentration was significantly increased in patients with advanced gastric cancer and can serve as a potential biomarker for chemotherapy monitoring[37]. Here we sought to investigate predictive value of cfDNA in efficacy of treatment and prognosis for NSCLC patients with chemotherapy, targeted therapy, immunotherapy or combined treatment.

## MATERIALS AND METHODS

### Study design and patient selection

This study is a single-institution protocol to evaluate peripheral cfDNA as a potential prognostic biomarker and efficacy predictor in NSCLC patients with chemotherapy or combination therapy. A total of 154 NSCLC patients who received chemotherapy or combined treatment in Jiangsu Cancer Hospital from December 2018 to February 2020 were enrolled. The clinical characteristics are shown in Table 1. Inclusion criteria include: 1) confirmed NSCLC diagnosis by pathohistology; 2) complete case data record. Exclusion criteria include: 1) patients with other malignant tumors; 2) patients with significant pre-existing cardiac, hepatic or renal disease; 3) patients with acute or chronic infectious disease; and 4) patients with mental illness prohibiting informed consent. All participants signed the informed consent agreement. The study was approved by the clinical research ethics committee of the Jiangsu Cancer Hospital and was conducted following the Declaration of Helsinki.

**Table 1.** Baseline Characteristics of Patients

| Characteristics | Number (N=154) | Proportion |
| --- | --- | --- |
| Sex & Age |  |  |
| Male | 107 | 69.5% |
| Female | 47 | 30.5% |
| Median age (yr) | 62 (34-79) |  |
| Histology |  |  |
| LUAD | 128 | 83.1% |
| LUSC | 26 | 16.9% |
| Clinical Stage |  |  |
| I | 2 | 1.3% |
| II | 4 | 2.6% |
| III | 19 | 12.3% |
| IV | 126 | 81.8% |
| ECOG (Eastern Cooperative Oncology Group, baseline) |  |  |
| 0 | 6 | 3.9% |
| 1 | 72 | 46.8% |
| 2 | 1 | 0.7% |
| NA | 75 | 48.7% |

### Assessment of peripheral cfDNA

All patients were subjected to peripheral blood samples collection before (baseline) and after (post-therapy) the first target cycle of chemotherapy. The cfDNA concentration was determined by QuantiDNA Direct cfDNA Test Kit (Diacarta. Inc., CA, USA) according to the manual and our previous publication[37]. In brief, 2-3 ml peripheral blood was drawn and subjected to 10 minutes centrifugation at 3000 rpm for plasma isolation. 50 μL of plasma was used for direct cfDNA detection according to manufacturer’s instructions.

### Efficacy and prognosis evaluation

The efficacy of treatment and prognosis were evaluated based on RECIST1.1 (Response Evaluation Criteria in Solid Tumors, Version 1.1)[38]. The criteria were as follows: Complete Response (CR): absence of all measurable lesions, or all residual lesions lower than diagnostic threshold (10 mm for the longer diameter of tumors and 15 mm for the shorter diameter of lymph nodes); Partial Response (PR): tumor burden (TB) reduced by >30% compared with baseline and the overall decrement ≥ 5 mm; Progressive Disease (PD): new measurable lesions or initial lesions increased by ≥ 20%; Stable Disease (SD): all which cannot be classified as CR, PR, or PD. Progression-free survival (PFS)[39] was the primary outcome that was defined as the days from the date of initial chemotherapy until the date of progressive disease, death, or the last follow up if progression or death had not occurred.

### Statistical analysis

We stratified the treatment evaluation by dug combination regiment which consisted of four groups: (1) chemotherapy only; (2) chemotherapy plus VEGF/VEGF receptor inhibitors (VEGFIs); (3) chemotherapy plus tyrosine kinase inhibitors (TKIs); and (4) chemotherapy plus immune checkpoint inhibitors (ICIs). The primary outcome was (1) progression-free survival (PFS); and secondary outcomes was (2) objective response ratio (ORR), defined as the proportion of CR and PR in all subjects. An initial model without interactions was used to identify the prognostic impact of baseline cfDNA, post-therapy cfDNA, and the cfDNA ratio respectively. Other demographic or clinical factors which may be associated with PFS were also evaluated via univariate Cox model separately and multivariate Cox model together. Survival curves were plotted by the Kaplan-Meier method with R package ‘survival’ and ‘survminer’.

## RESULTS

### Pathological and demographic characteristics

The pathological and demographic characteristics of the 154 patients were summarized (**Table1**). The median age was 62 years (34-79); 107 (69%) of participants were male and 47 (31%) were female. Since our aim is evaluating cfDNA clinical utilization, the most typical patients (LUAD in late stage) were selected. For example, 128 (83%) of patients were LUAD, and 26 (17%) were LUSC. 126 (82%) were in stage IV, 19 (12%) were in stage III and others were in stage I/II.

### Peripheral cfDNA baseline correlates with tumor burden

To assess the relationship between cfDNA and TB, we defined TB_baseline as the pre-treatment TB, and only selected those whose interval between cfDNA test and TB evaluation was within 7 days (N=80). We defined TB_post-chemotherapy as the post-chemotherapy TB and restricted the interval between cfDNA test and TB evaluation to no more than 7 days (N=47). Overall, a weakly positive correlation between TB and cfDNA was observed at baseline (*N*=80, Pearson’s coefficient = 0.24; 95% confidence interval (CI): 0.017-0.433; *P* = 0.03, **Fig 1A**), while no significant correlation was found for post-chemotherapy (*N*=47, Pearson’s coefficient = 0.124; 95% CI: −0.169-0.397; *P* = 0.4, **Fig 1B**).

**Fig 1.**
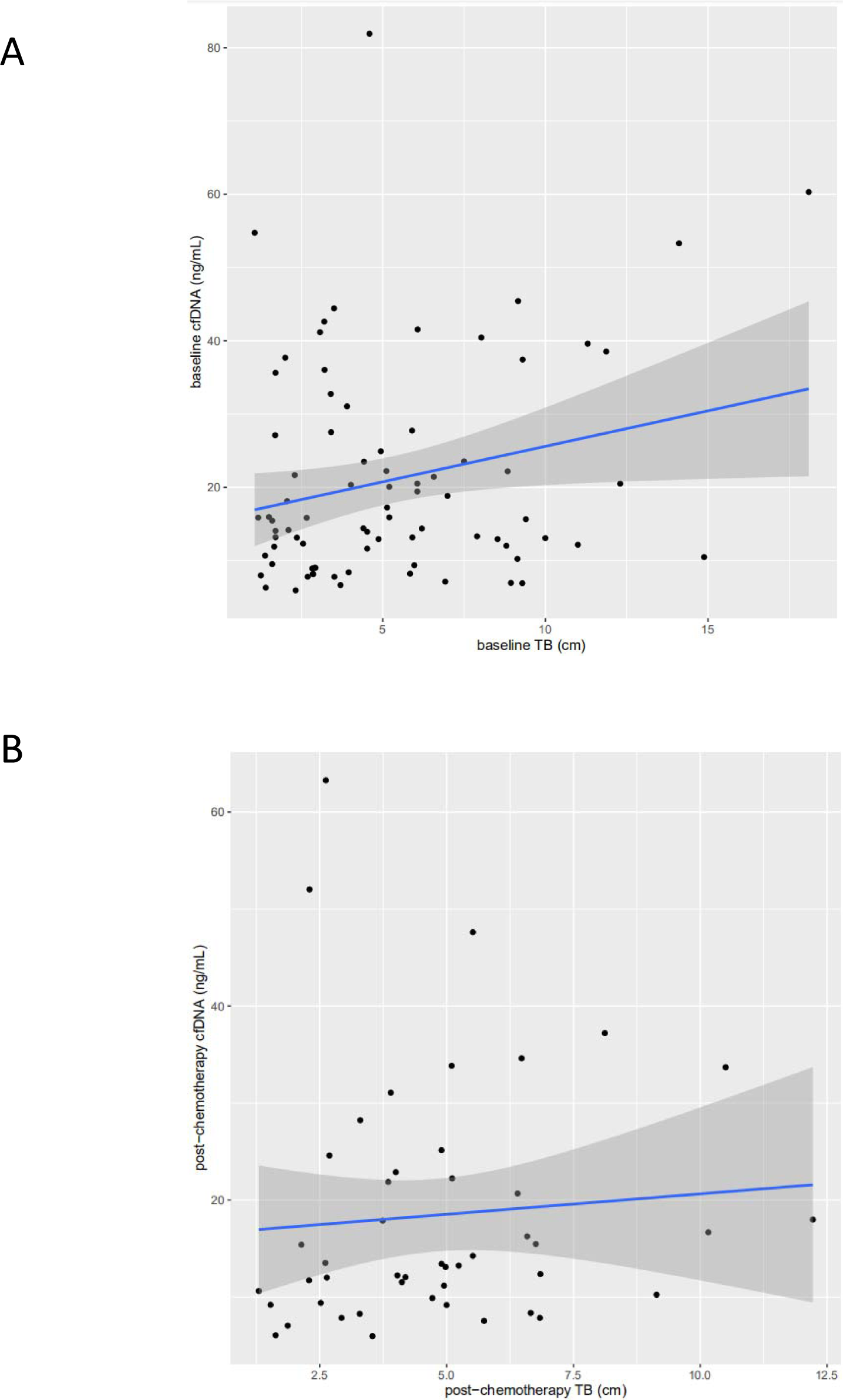
Scatter plot showing a weakly positive correlation of baseline cfDNA with baseline tumor burden. Tumor burden was evaluated by Response Evaluation Criteria in Solid Tumors, version 1.1. cfDNA was quantified by QuantiDNA Direct cfDNA Test Kit (Diacarta. Inc., CA, USA) according to the manual both (**A**) at baseline and (**B**) post-chemotherapy. We selected those whose interval between cfDNA test and TB evaluation was within 7 days, so 80 cases were qualified (A) at baseline, and 47 cases were qualified (B) post-chemotherapy. A weakly positive correlation between TB and cfDNA was observed at baseline (*N*=80, Pearson’s coefficient = 0.24; 95% CI: 0.017-0.433; *P* = 0.03), while no significant correlation was found for post-chemotherapy (*N*=47, Pearson’s coefficient = 0.124; 95% CI: −0.169-0.397; *P* = 0.4).

In addition, we also assessed other clinical factors which may be correlated with cfDNA. No significant correlations were found between age and cfDNA either at baseline (P=0.1) or post-chemotherapy (P=0.4), stage at baseline (P=0.9) or post-chemotherapy (P=0.4), ECOG score at baseline (P=0.8) or post-chemotherapy (P=0.8), gender at baseline (Wilcoxon rank sum test, P=0.5) or post-chemotherapy (Wilcoxon rank sum test, P=0.4). We also found no significant difference of cfDNA between LUAD and LUSC at baseline (Wilcoxon rank sum test, P=0.16) or post-chemotherapy (Wilcoxon rank sum test, P=0.4), or among different therapy regimens (**Supplementary Table 1**).

### Plasma cfDNA relates to objective response rate (ORR) and progression-free survival (PFS)

Since tumor burden usually correlates with clinical outcomes, we then investigated the relationship between clinical outcomes and peripheral cfDNA, we monitored peripheral cfDNA of all available patients (N=154) at baseline (121/154, 79% before chemotherapy by 0-7 days), post-chemotherapy (137/154, 89% after chemotherapy by 20-30 days) and derived Ratio (post-chemotherapy/baseline) for each patient.

Firstly, we compared the baseline cfDNA and post-chemotherapy cfDNA between responsive group (PR/CR, *N*=56) and non-responsive group (SD/PD, *N*=80). Overall, the responsive group trended toward higher baseline cfDNA (median 17.68 ng/mL) than the non-responsive (median 13.70 ng/mL) (*P*=0.058, Wilcoxon rank-sum test, **Fig 2A**). However, we found no significant difference in post-chemotherapy cfDNA between the two (*P*=0.6, Wilcoxon rank-sum test, **Fig 2B**), although the median post-chemotherapy cfDNA in the responsive (17.18 ng/mL) was modestly lower than that of the non-responsive (19.15 ng/mL). Notably we found a significantly lower ratio in the responsive group (median 0.87) than that of the non-responsive (median 1.21) (*P*=0.012, Wilcoxon rank-sum test, **Fig 2C**). These data suggested that cfDNA can be used to discriminate responsive patients from the non-responsive well, especially with cfDNA ratio which reflected the dynamic change of plasma cfDNA.

**Fig 2.**
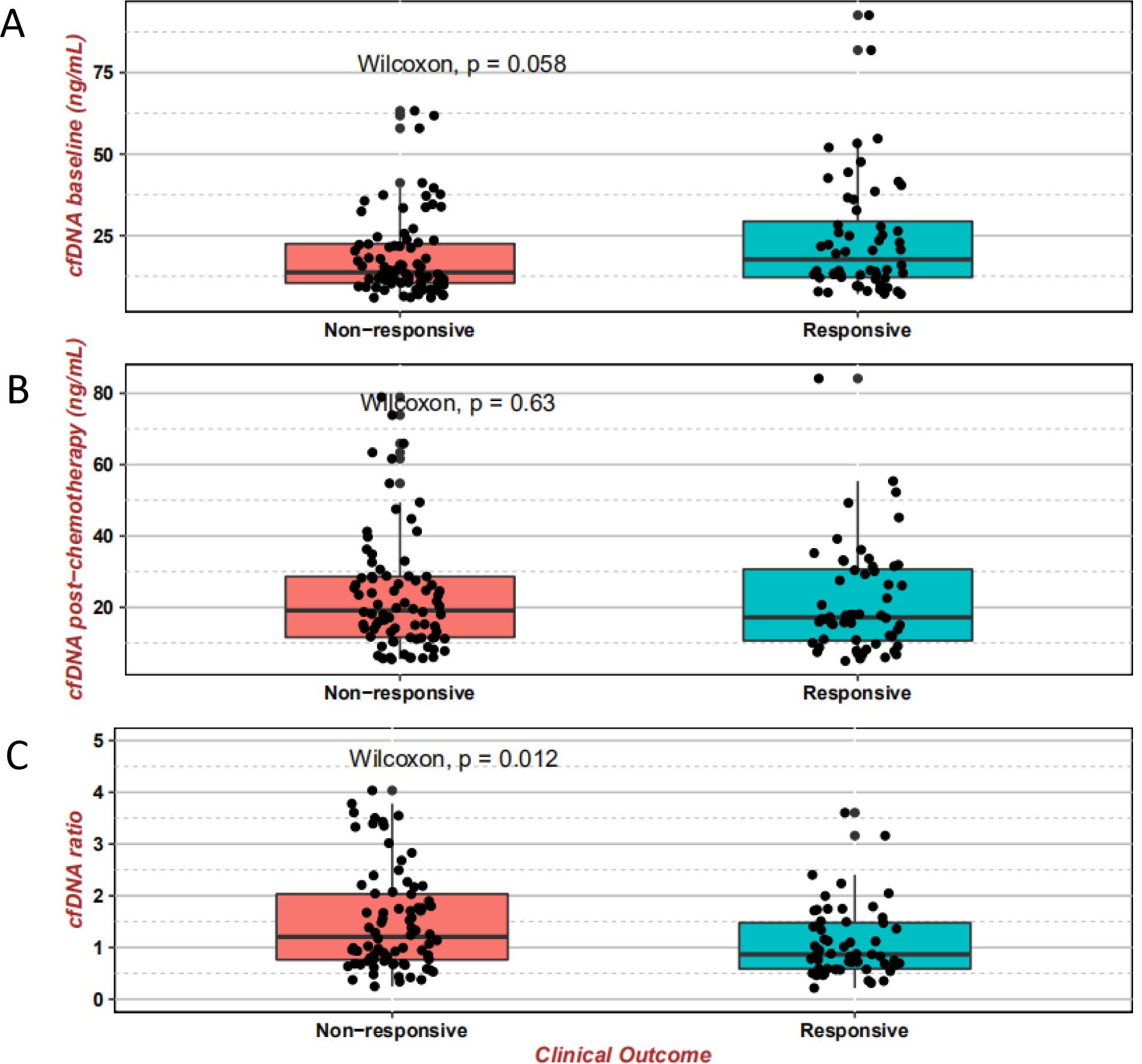
Comparison of cfDNA levels and cfDNA ratio between the responsive group and non-responsive group. Boxplots from top to bottom showed the baseline value (**A**), post-therapy value (**B**), and ratio value (**C**) of cfDNA respectively in both the responsive (PR/CR, *N*=56) group and non-responsive (PD/SD, *N*=80) group, the significance of difference between the two was estimated by Wilcoxon test.

Then we divided this cohort into Ratio_low and Ratio_high group by the median Ratio (1.0271). Similar comparative analysis was also carried out between Baseline_low and Baseline_high group (cut-value: median of cfDNA baseline, 15.43 ng/mL), and Post-chemotherapy_low and Post-chemotherapy_high group (cut-value:median of post-chemotherapy cfDNA, 18.42 ng/mL), respectively.

Significantly improved PFS benefit was observed for Ratio_low (HR: 0.54 (95% CI: 0.29-1.01); Log-rank test, *P*=0.05, **Fig 3A**) compared with Ratio_high, while no significant difference was found between Baseline_low and Baseline_high group (Log-rank test, *P*=0.86, **Fig 3B**) and between Post-chemotherapy_low and Post-chemotherapy_high group (Log-rank test, *P*=0.57, **Fig 3C**). After a median follow-up of 6.4 months, the median PFS of Ratio_low group was 6.1 months which was 2 months longer than that of Ratio_high group (4.1 months). The objective response ratio (ORR) of the Ratio_low group (33/77, 42.8%) was also 1.5 times higher than that of the Ratio_high group (22/77, 28.5%).

**Fig 3.**
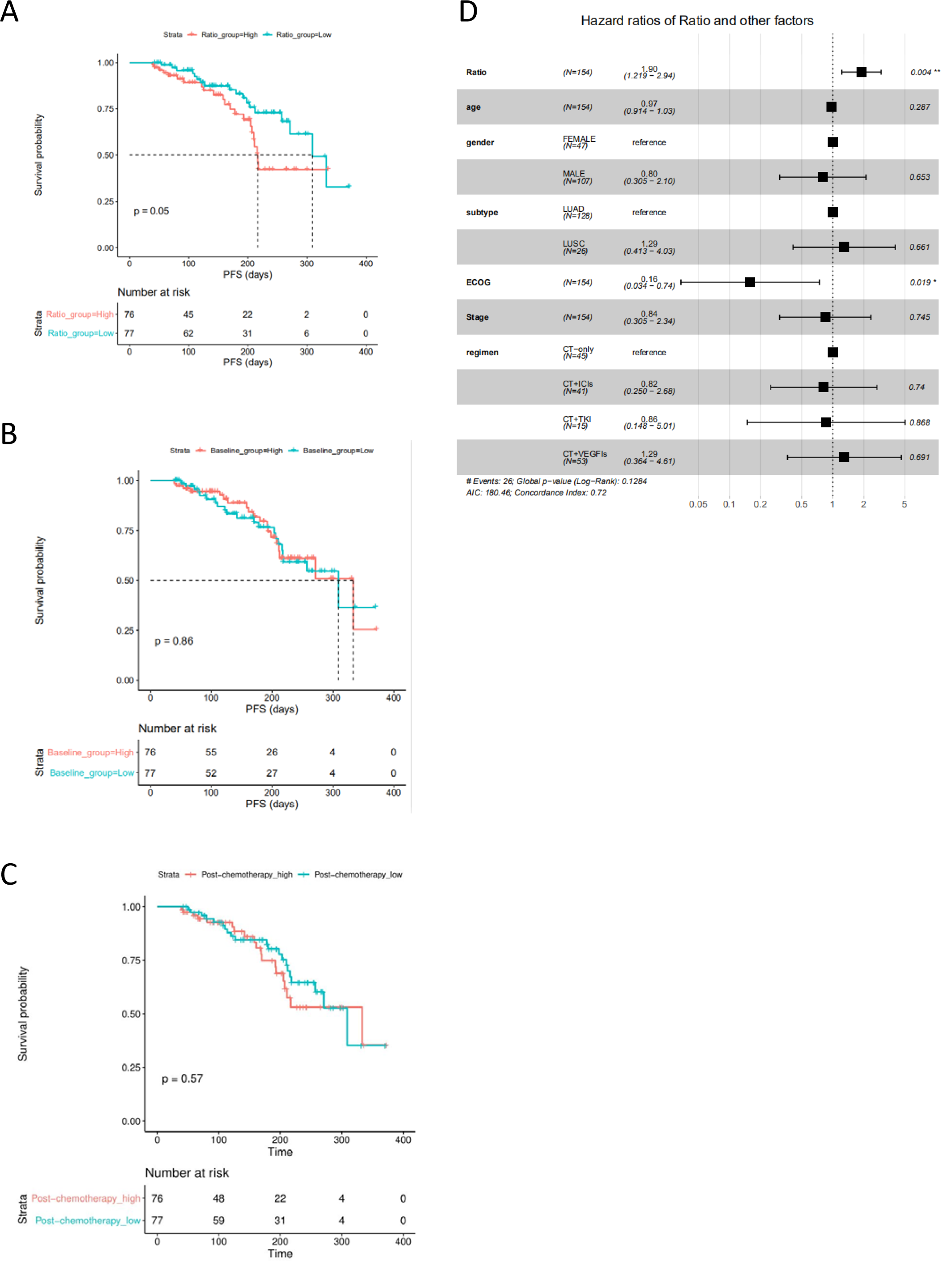
Progression-free Survival in the overall cohort (*N*=154). Kaplan-Meier curves for comparisons of progression-free survival between **(A)** high cfDNA Ratio and low cfDNA Ratio groups, **(B)** high cfDNA baseline and low cfDNA baseline groups, **(C)** high post-therapy cfDNA and low post-therapy cfDNA groups (cut-values were set as median value), respectively. **(D)** The hazard ratios of cfDNA ratio and other important clinical factors by multivariate Cox model. Cut-values were set as the median value of the overall cohort, respectively. Progression-free survival was assessed according to Response Evaluation Criteria in Solid Tumors, version 1.1 through investigators’ review, and tick marks represent data censored at the last time the patient was known to be alive and without disease progression.

Other factors which may impact PFS were evaluated by univariate Cox model such as age (HR:1.00 (95% CI: 0.96-1.03); *P*=0.8), gender (HR:1.01 (95% CI: 0.51-2.00); *P*=1.0), subtype (LUSC v.s. LUAD, HR:1.36 (95% CI: 0.64-2.86); *P*=0.4), ECOG (HR:0.29 (95% CI: 0.08-1.00); *P*=0.05), stage (HR:0.86 (95% CI: 0.42-1.76); *P*=0.7), therapy regimen (chemotherapy+ICIs v.s. chemotherapy, HR:0.77 (95% CI:0.33-1.78); *P*=0.5), chemotherapy+TKIs v.s. chemotherapy, HR:0.69 (95% CI:0.23-1.97); *P*=0.48), chemotherapy+VEGFIs v.s. chemotherapy, HR:0.68 (95% CI: 0.30-1.52); *P*=0.35). None of these factors showed a significant impact on PFS by univariate or multivariate Cox model in our study (**Fig 3D**).

To exclude potential effects of predefined low/high groups, we also evaluated the correlation between cfDNA and PFS by both univariate Cox model and multivariate Cox model. Similar log Log-rank test results, only cfDNA ratio (HR: 1.55 (95% CI: 1.11-2.18); *P*=0.01) was significantly negatively related with PFS in univariate Cox model, but not cfDNA baseline (HR: 0.99 (95% CI: 0.97-1.01); *P*=0.4) and post-therapy cfDNA (HR: 1.01 (95% CI: 0.99-1.03); *P*=0.4). While In multivariate Cox model (taking into account age, gender, subtype, ECOG, stage, and therapy regimen), both cfDNA baseline (HR:0.95 (95% CI: 0.91-1.00); *P*=0.03) and cfDNA ratio (HR:1.90 (95% CI: 1.2-2.95); *P*=0.004) were shown to be significantly related with PFS, but not post-therapy cfDNA (HR:1.02 (95% CI: 0.99-1.05); *P*=0.28).

Furthermore, we compared the demographic (age and gender), pathological (subtype, stage, and ECOG scores), and therapeutic (therapy regimens) characteristics between Ratio_low and Ratio_high group, and found no significant difference (chi-square test) in all these factors (**Table 2**), which indicated the evenly distributed patient between these two groups.

**Table 2.** Comparisons between Ratio_high group and Ratio_low group

| Ratio | High | Low | P |
| --- | --- | --- | --- |
| n | 77 | 77 |  |
| age (mean (SD)) | 60.30 (8.68) | 61.65 (9.16) | 0.349 |
| gender = MALE (%) | 53 (68.8) | 54 (70.1) | 1 |
| subtype = LUSC (%) | 9 (11.7) | 17 (22.1) | 0.132 |
| ECOG (mean (SD)) | 0.95 (0.32) | 0.93 (0.27) | 0.721 |
| Stage (mean (SD)) | 3.75 (0.59) | 3.81 (0.51) | 0.483 |
| response (%) |  |  | 0.157 |
| PD | 17 (22.4) | 11 (14.3) |  |
| PR | 22 (28.9) | 33 (42.9) |  |
| SD | 37 (48.7) | 33 (42.9) |  |
| PFS/days (mean (SD)) | 139.59 (76.16) | 179.08 (83.75) | 0.003 |
| regimen (%) |  |  | 0.227 |
| CT-only | 25 (32.5) | 20 (26.0) |  |
| CT+ICIs | 17 (22.1) | 24 (31.2) |  |
| CT+TKIs | 5 (6.5) | 10 (13.0) |  |
| CT+VEGFIs | 30 (39.0) | 23 (29.9) |  |

### Stratification analysis by subtype

Since LUAD and LUSC are two main pathologic subtypes of NSCLC with different clinical managements and prognostics, we further analyzed the prognostic significance of the cfDNA baseline, cfDNA post-therapy, and cfDNA ratio in these two subgroups, respectively.

For the LUAD group (*N*=128), we found a significantly improved PFS benefit for the Ratio_low group (HR: 0.42 (95% CI: 0.20-0.86); *P*=0.015, **Fig 4A**) compared with Ratio_high group. The median PFS of Ratio_low group was 6.3 months which was 2.1 months longer than that of Ratio_high group (4.2 months). Additionally, ORR of Ratio_low group (43.3%) was also higher than that of the Ratio_high group (29.4%).

**Fig 4.**
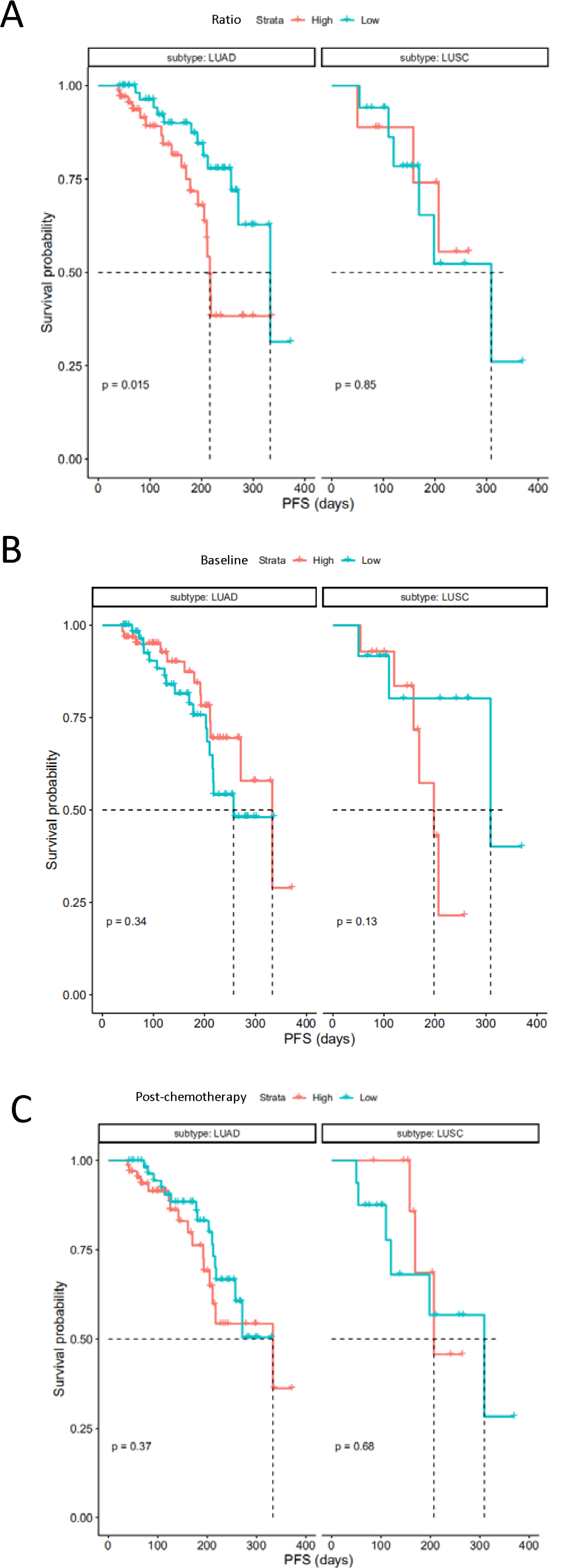
Progression-free Survival pathological subtype. Stratification analysis of progression-free survival by pathological subtype (LUAD, *N*=128 and LUSC, *N*=26) **(A)** high cfDNA Ratio and low cfDNA Ratio groups, **(B)** high cfDNA baseline and low cfDNA baseline groups, **(C)** high post-therapy cfDNA and low post-therapy cfDNA groups (cut-values were set as median value), respectively. Progression-free survival was assessed according to Response Evaluation Criteria in Solid Tumors, version 1.1 through investigators’ review, and tick marks represent data censored at the last time the patient was known to be alive and without disease progression.

The LUSC group, as a minority of our cohort (*N*=26), no significantly improved PFS benefit was found for the Ratio_low group (HR: 1.12 (95% CI: 0.27-4.91); *P*=0.85, **Fig 4A**) compared with Ratio_high group. With only 2 patients followed past 300 days, both in the Ratio low group, the median PFS was 4.9 months in the Ratio low compared to 6.8 months in the Ratio_high group. However, ORR of Ratio_low group (41.1%) was trended higher than that of the Ratio_high group (22.2%).

Similar to the whole cohort, no significant difference of PFS was found between Baseline_low and Baseline_high group (**Fig 4B**) or between Post-chemotherapy_low and Post-chemotherapy_high group (**Fig 4C**) when stratified by LUAD and LUSC, respectively.

### Stratification analysis by treatment

Since the change of cfDNA (Ratio) during treatment strongly correlated with PFS and objective response as shown above, we further analyzed in 4 subgroups stratified by therapy regimen: (1) chemotherapy only; (2) chemotherapy plus VEGF/VEGF receptor inhibitors (VEGFIs); (3) chemotherapy plus TKIs; (4) chemotherapy plus ICIs.

Only the Ratio_low group of patients received chemotherapy plus VEGFIs treatment showed significantly prolonged PFS compared to those in Ratio_high group (HR: 0.23, 95% CI: 0.06-0.88; *P*=0.02, **Fig 5A**). Additionally, ORR of Ratio_low group (9/23, 39%) was numerically higher than that of Ratio_high group (8/30, 27%). Importantly, only 2 (8.6%) of Ratio_low had Progressive Disease (PD), while 7 (23.3%) of Ratio_high had PD. In other therapy groups, no significant difference of PFS was found, e.g., chemotherapy-only group (HR=0.82, *P*=0.7), chemotherapy with TKIs group (HR=0.68, *P*=0.7), chemotherapy with ICIs group (HR=0.72, *P*=0.6). Whereas, ORR of Ratio_low group with all these three regimens are numerically higher than that of Ratio_high group (40% vs 28% for chemotherapy-only, 50% vs 20% for chemotherapy with TKIs, 46% vs 35% for chemotherapy with ICIs).

**Fig 5.**
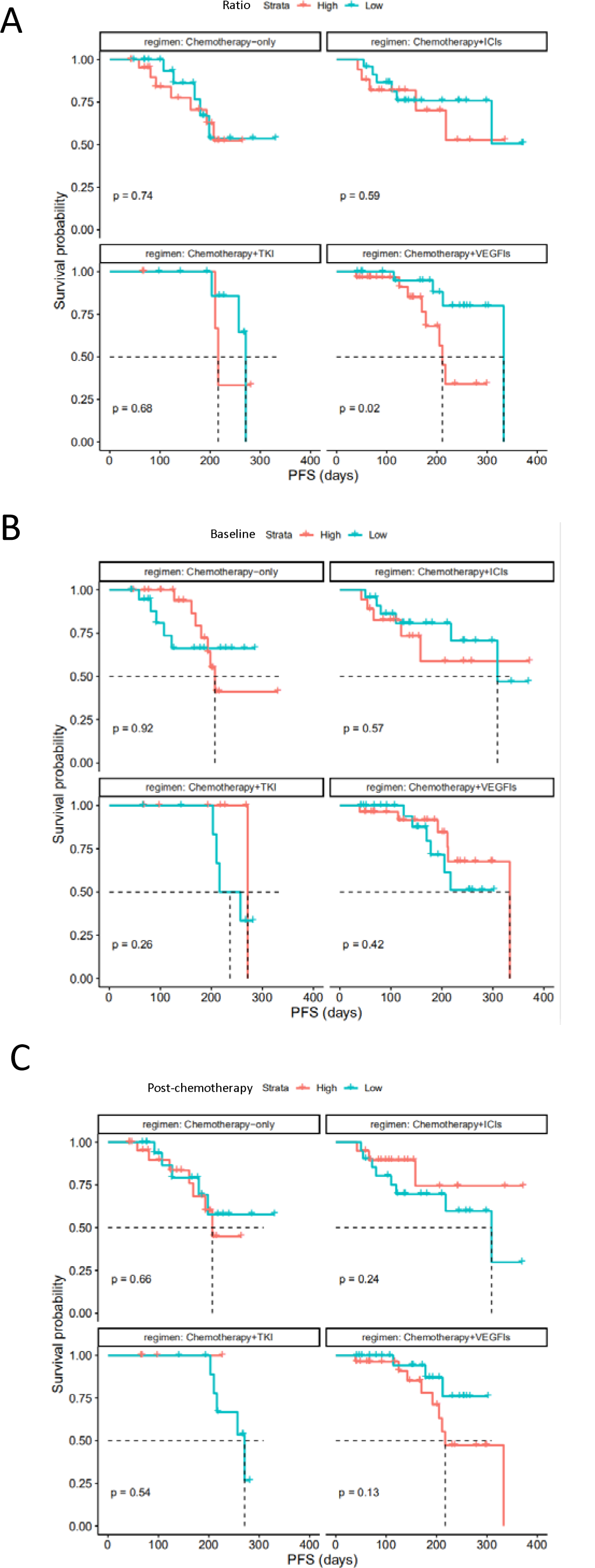
Progression-free Survival in subgroups by therapy regimen. Stratified analysis of Kaplan-Meier curves by therapy regimen (chemotherapy alone (*N*=45), chemotherapy plus TKIs (*N*=15), chemotherapy plus VEGFIs (*N*=53), and chemotherapy plus ICIs (*N*=41)) for comparisons of progression-free survival between **(A)** high cfDNA Ratio and low cfDNA Ratio groups, **(B)** high cfDNA baseline and low cfDNA baseline groups, **(C)** high post-therapy cfDNA and low post-therapy cfDNA groups (cut-values were set as median value), respectively. Progression-free survival was assessed according to Response Evaluation Criteria in Solid Tumors, version 1.1 through investigators’ review, and tick marks represent data censored at the last time the patient was known to be alive and without disease progression.

In addition, no significant difference of PFS was found between Baseline_low and Baseline_high group (**Fig 5B**) or Post-chemotherapy_low and Post-chemotherapy_high group (**Fig 5C**) when stratified by these therapy regimens, respectively.

## DISCUSSION

Over the past two decades, important advancements have been achieved in the treatment of advanced NSCLC with our increasing understanding of the disease biology, tumorigenesis, early detection and multimodal care[2]. Notably, the utilization of targeted therapy and immunotherapy has led to remarkable survival benefits in selected patients[7, 8]. However, there is no universal, reliable biomarker to predict or evaluate the treatment response and prognosis of different managements. In this study, we confirmed that the kinetics of plasma cfDNA (Ratio, post-chemotherapy/baseline) is well correlated with clinical response (ORR) and progression free survival (PFS) in chemotherapy with VEGF inhibitor targeted therapy, functioning as a potential, convenient predictive biomarker for NSCLC patients.

Circulating cfDNA is derived from a combination of apoptosis, necrosis and active secretion from both cancer cells and normal cells which are subjected to harsh stimuli such as chemotherapy[40] or driven by inflammatory process[41]. It was found at higher levels in patients with advanced cancer than in early stage disease or healthy individuals[42, 43]. In the present study, we found a positive correlation between tumor burden and cfDNA baseline in NSCLC (Fig 1). Although the tumor-derived fraction of these total cfDNA (ctDNA) has been widely investigated as a prognostic biomarker in various cancer types including breast, colon and lung cancer[18, 44-46], the main challenges are low amount of ctDNA, detection cost and reproducibility limitations. For example, some typical difficulties of NGS application in this scenario include inadequate analytical sensitivity and specificity, such as detection limit of low allelic frequencies, and sequencing false positive[43, 47]. The total cfDNA with its higher feasibility has become an attractive alternative biomarker[15-17]. Our study utilized proven fluorescent probes and quick turnaround time of the SuperbDNA™ technology to measure plasma total cfDNA[37, 48]. Indeed, the Ratio (post-chemotherapy/baseline) of cfDNA show a correlated with clinical treatment response. The responsive patients obviously have much lower Ratio than those with no response individuals (Fig 2). In addition, the post-chemo/base ratio, but not baseline and post-chemotherapy, levels of cfDNA has a reversed correlation with progression-free survival (PFS) evaluated by RECIST (response evaluation criteria in solid tumors) criteria (p=0.05), combining all cases regardless of the therapy regimens (chemotherapy only, chemotherapy plus targeted therapy or immunotherapy) they received (Fig 3). With Ratio cutoff-value set at either median (1.03), Ratio_low group has a significantly improved PFS with 2 months longer than that of Ratio_high group (4.1 months) (Fig 3). Unlike other studies showing a correlation between a single snapshot of elevated cfDNA concentration and poor survival[33, 36, 49], our data revealed that the response of cfDNA is an effective response indicator. To avoid potential effects of predefined cutoff-value of low/high Ratio, we also performed the correlations of PFS with each individual cfDNA baseline, post-chemotherapy and Ratio. Similarly, cfDNA Ratio, but not baseline or post-chemotherapy, was significantly negatively related with PFS (*P*=0.01). Interestingly, when stratified by pathohistology, the predicted value of the cfDNA Ratio was only significant in the LUAD group. The LUSC group was smaller, with only 2 patients exhibiting tumor progression. Larger studies are needed to determine the utility of the cfDNA Ratio in LUSC patients. Among different therapy regimens, the strong negative correlation between PFS and Ratio was reproduced in the patient with chemotherapy plus VEGFIs (P=0.02), but not chemotherapy only (P=0.74), chemotherapy plus TKI (P=0.68) or immunotherapy (P=0.59). It may attribute to a relative short term of follow-up, insufficient case number or non-molecular preselection, since reports have shown that targeted therapy (TKI) mostly benefits NSCLC patients with driver (such as EGFR) mutations[50, 51] and immunotherapy usually takes a longer time to be clinically effective[52]. Further study targeting molecularly selected patients with a larger scale and longer follow-up is needed for validations.

In terms of clinical treatment response, objective response rate (ORR) showed the similar patterns as PFS. Only cfDNA Ratio, not baseline or post-chemotherapy, distinguished the subgroups who had a better clinical response and beneficial outcomes. Our results are consistent with the previous report that monitoring of plasma DNA change during chemotherapy identifies patients who are likely to exhibit a therapeutic response or disease progression[53]. Other studies however suggested that cfDNA concentration is not reliable enough to predict treatment response in NSCLC when treatment is chemotherapy[36, 54]. One possible explanation is that sensitivity to chemotherapy and cfDNA levels during treatment may vary among individuals, or depend on timing of the sample acquisition[34]. We selected evaluating cfDNA level after one cycle of chemotherapy (post treatment 20-30 days) based on the consideration that cfDNA would remain relatively stable during cycles and early evaluation could allow for therapeutic adjustment if needed. PFS can be measured but the results are too late to allow for therapeutic modification. ORR is a quicker index but still requires an imaging cycle and detailed image evaluation. The cfDNA Ratio is measured after the first cycle and is immediately interpretable, allowing for real time treatment adjustments. The Ratio_low group enjoyed an ORR more than 1.5 times higher than that of Ratio_high group (42.8% vs 28.5%) regardless of treatment regimen. The effect was most pronounced in the chemotherapy plus VEGFIs group, only % of patients in Ratio_low group had disease progressed (PD), while in Ratio_high group the number increased to 23.3%. These data support the predictive role of cfDNA ratio in efficacy of chemotherapy.

To our knowledge, this is the first study to utilize the concept of cfDNA ratio (post-chemotherapy/baseline) to better present personalized medicine management. The concentration of cfDNA varies among individuals based on personalized nuances of the physiology and tumor characteristics[55]. Using a cfDNA ratio, captured in an appropriate time interval, normalizes the physiological effects leading to an estimate of tumor response. Indeed, from our data, the snapshot of baseline cfDNA did correlate with some clinical parameters like tumor burden. Yet for clinical response (ORR) and prognostic prediction, the cfDNA ratio seems to provide an early measure of tumor response.

This first study has limitations. (1) neither ORR or PFS are fully predictive of overall survival (OS). However, both ORR and PFS are used clinically to alter therapy, and an even earlier measure of tumor response would be advantageous. (2) Basal release and accumulation of cfDNA in the plasma, as mentioned above, is not an identical for every tumor or every patient. We have provided data supporting its imperfect potential for measuring basal TB in gastric cancer[37] and now in NSCLC. S will other common tumor markers (ex. CEA, PSA), more tumor subtypes should be screened for further validations. (3) cfDNA is quick, accurate, and inexpensive, but it is not specific for cancer. Logically ctDNA or other tests, if they can be made quantitative and reliable would be useful as adjuncts to calibrate the cfDNA test.

As with most initial discoveries, this is a single institution study that promises to advance a simple test that can provide an early indicator of NSCLC response to a number of different systemic therapies. We believe it should advance to a larger, multi-center trial.

---

**Supplementary Table 1.**
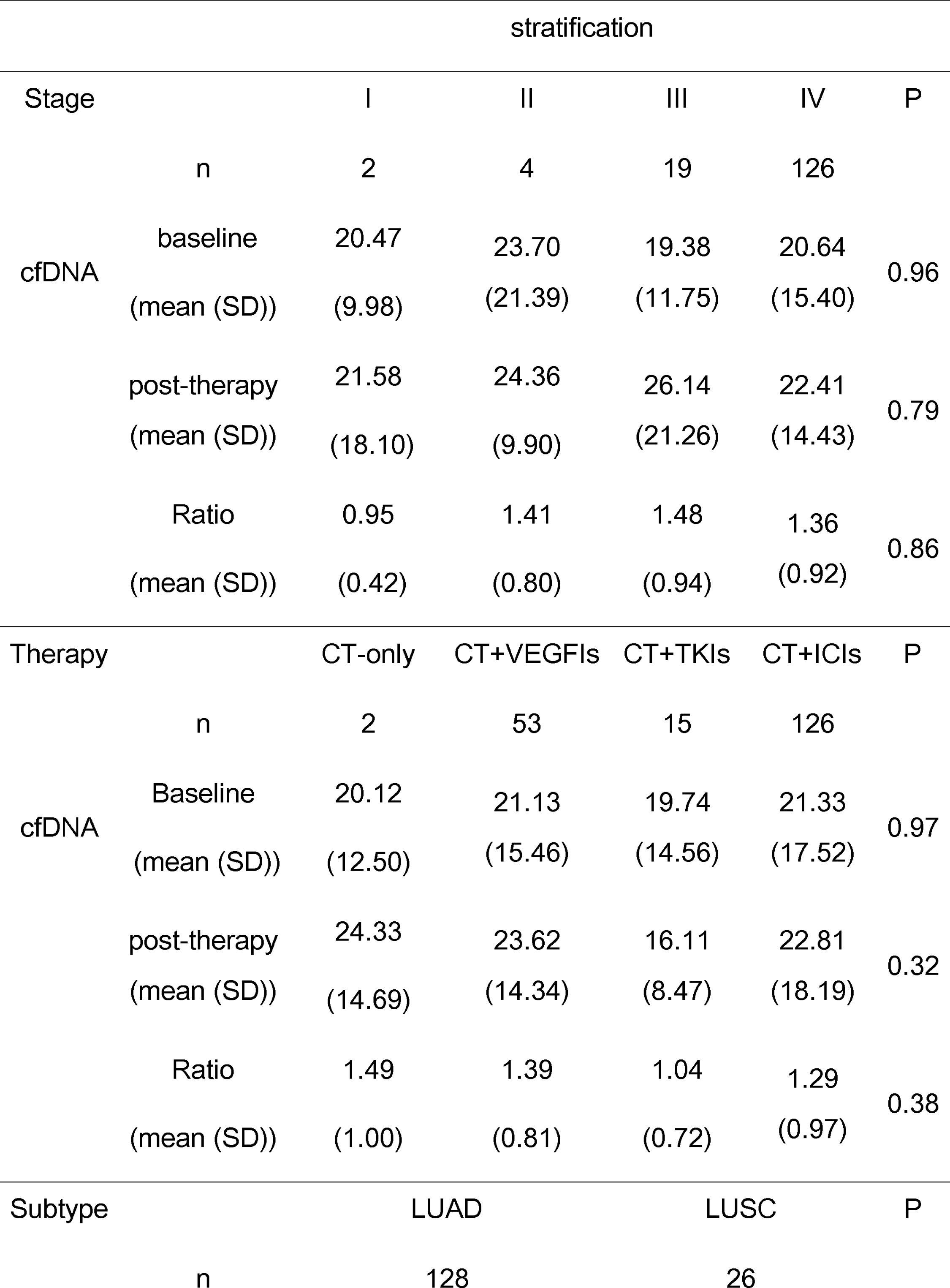
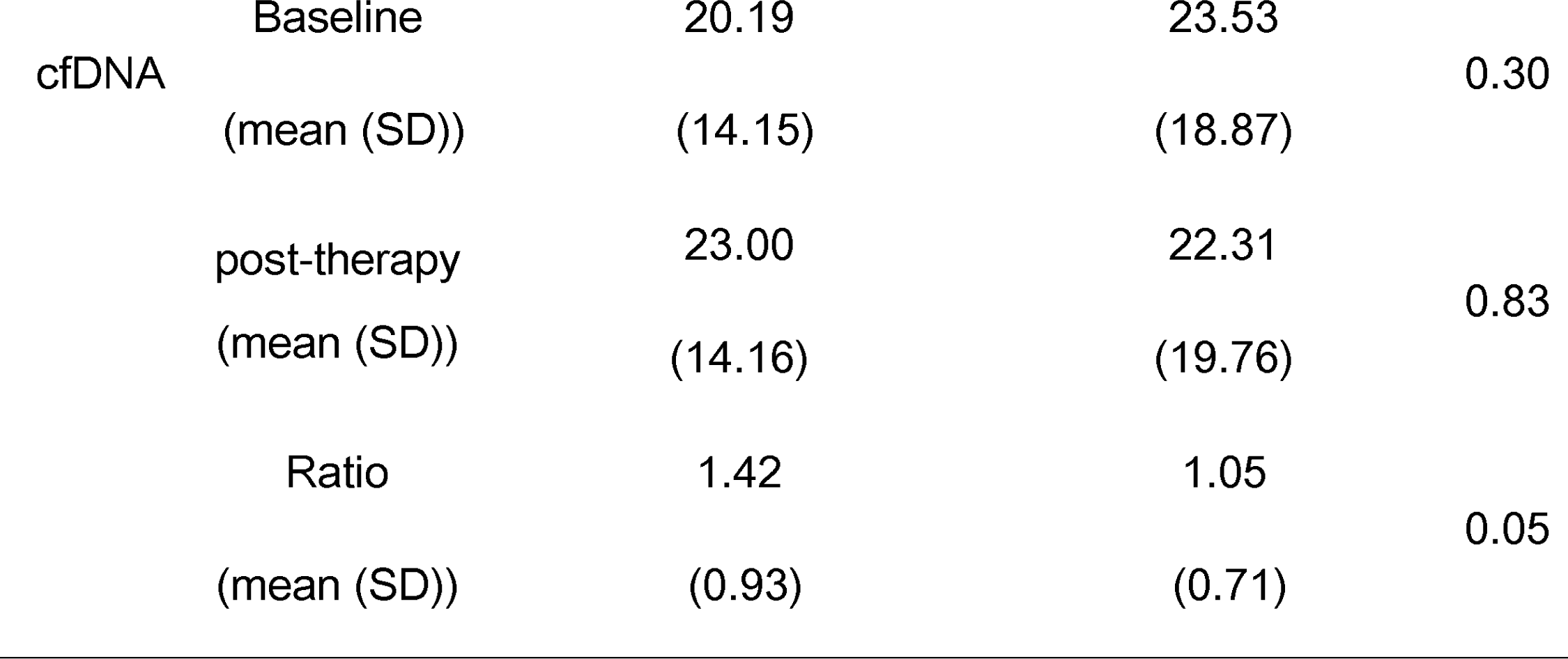
Comparisons of cfDNA levels and ratio by stage, therapy, and subtype.

Comparisons of cfDNA levels and ratio by stage, therapy, and subtype
| stratification |  |  |  |  |  |  |
| --- | --- | --- | --- | --- | --- | --- |
| Stage |  | I | II | III | IV | P |
|  | n | 2 | 4 | 19 | 126 |  |
| cfDNA | baseline | 20.47 | 23.70 | 19.38 | 20.64 | 0.96 |
|  | (mean (SD)) | (9.98) | (21.39) | (11.75) | (15.40) |  |
|  | post-therapy | 21.58 | 24.36 | 26.14 | 22.41 | 0.79 |
|  | (mean (SD)) | (18.10) | (9.90) | (21.26) | (14.43) |  |
|  | Ratio | 0.95 | 1.41 | 1.48 | 1.36 | 0.86 |
|  | (mean (SD)) | (0.42) | (0.80) | (0.94) | (0.92) |  |
| Therapy |  | CT-only | CT+VEGFIs | CT+TKIs | CT+ICIs | P |
|  | n | 2 | 53 | 15 | 126 |  |
| cfDNA | Baseline | 20.12 | 21.13 | 19.74 | 21.33 | 0.97 |
|  | (mean (SD)) | (12.50) | (15.46) | (14.56) | (17.52) |  |
|  | post-therapy | 24.33 | 23.62 | 16.11 | 22.81 | 0.32 |
|  | (mean (SD)) | (14.69) | (14.34) | (8.47) | (18.19) |  |
|  | Ratio | 1.49 | 1.39 | 1.04 | 1.29 | 0.38 |
|  | (mean (SD)) | (1.00) | (0.81) | (0.72) | (0.97) |  |
| Subtype |  | LUAD |  | LUSC |  | P |
|  | n | 128 |  | 26 |  |  |

|  |  |  |  |  |
| --- | --- | --- | --- | --- |
| cfDNA | Baseline | 20.19 | 23.53 | 0.30 |
|  | (mean (SD)) | (14.15) | (18.87) |  |
|  | post-therapy | 23.00 | 22.31 | 0.83 |
|  | (mean (SD)) | (14.16) | (19.76) |  |
|  | Ratio | 1.42 | 1.05 | 0.05 |
|  | (mean (SD)) | (0.93) | (0.71) |  |

## Notes

### Competing Interest Statement

The authors have declared no competing interest.

